# Production of Fc-fused receptor agonists for glucagon-like peptide-1/glucose-dependent insulinotropic polypeptide (GLP-1/GIP) in the milk of transgenic mice

**DOI:** 10.1101/2025.03.17.643593

**Authors:** Yu Rao, Shuai Yu, Bao-Zhu Wang, Sheng Cui, Ke-Mian Gou

## Abstract

Incretin hormones, including glucagon-like peptide-1 (GLP-1) and glucose-dependent insulinotropic polypeptide (GIP), play pivotal roles in glucose homeostasis and metabolic regulation. Therapeutic incretin receptor agonists (RAs), such as tirzepatide, are widely used to manage type 2 diabetes and obesity. However, incretin RAs are facing production challenges at present. Therefore, we engineered transgenic (tg) mice to secrete incretin RAs in milk, leveraging mammary gland bioreactors for cost-effective peptide production. The goat beta-casein promoter-driven constructs encoding tirzepatide-derived peptide linked to human IgG4 Fc via a (GGGGS)_3_ spacer were used to produce tg mice. Founders tg-1 and tg-5 exhibited mammary-specific expression, yielding 0.8–1.42 g/L recombinant protein exclusively in milk. Progeny nursed by founders showed sustained hypoglycemia (10−39% reduction; p < 0.05) and marked weight loss (14−49%; p < 0.01) compared to wild-type controls, validating the bioactivity of milk-derived GLP-1/GIP RAs. Moreover, tg-5-nursed offspring experienced high mortality post-Day 16, likely due to overdosing. This proof-of-concept demonstrates the mammary gland bioreactor as a viable platform for incretin RAs production, circumventing complex synthesis and enabling scalable biologics manufacturing.

## 1. Introduction

Incretin hormones encompass glucagon-like peptide-1 (GLP-1) and glucose-dependent insulinotropic polypeptide (GIP). Two GLP-1 receptor agonists (RAs), Dulaglutide ^1^ and Semaglutide ^2^, are used primarily for treating type 2 diabetes and obesity, with extended benefits to cardiovascular and/or chronic kidney disease ^3^. The GLP-1 protein shows a highly conservative receptor-binding domain (residues 7-37) comprising the sequence HAEGTFTSDVSSYLEGQAAKEFIAWLVKGR, which the current therapeutic GLP-1 RAs are fully chemically synthesised or semi-biosynthesised based on this peptide. During this process, sequence optimisation is routinely implemented to enhance pharmacokinetics, exemplified by the substitution of Ala-8 to Gly-8 in GlaxoSmithKline’s Tanzeum or to alpha-aminoisobutyric acid (al-pha-Aib) in Novo Nordisk’s Semaglutide to resist dipeptidyl peptidase-4 degradation. Moreover, a long-chain fatty acid is also usually conjugated at Lys-26, as in Semaglutide, to augment the binding of RAs to albumin for prolonged plasma half-life.

On the other hand, Fc-fusion technology which conjugates bioactive peptides to the immunoglobulin Fc domain can extend plasma half-life and simplify the purification of aimed peptides ^4^. It has been successfully demonstrated that Dulaglutide, an IgG4 Fc-fused GLP-1 RA, extends plasma half-life by approximately one week and can efficiently improve glycaemic control for patients with type 2 diabetes ^5^.

GIP exhibits glucagon suppression during hyperglycaemia and glucagonotropic activity in hypoglycaemia, offering prophylactic benefits against hypoglycaemia. Recent investigation has demonstrated that long-acting GIP RAs depend crucially on GIP receptor signalling in inhibitory GABAergic neurons to decrease body weight and food intake in mice ^6^. Further study into dual GLP-1/GIP co-agonism revealed synergistic enhancement of β-cell insulin secretion, insulin sensitivity improvement, glucose-dependent glycaemic regulation, lipid-lowering, and profound weight loss ^7^. This pharmacological synergy underpinned the development of Tirzepatide.

Tirzepatide is designed to activate GIP and GLP-1 receptors simultaneously to enhance insulin secretion in response to nutrient intake, and this dual action helps to improve glycemic control and promote weight loss ^8^. Tirzepatide incorporates a 39-residue sequence (Y(alpha-Aib)EGTFTSDYSI(alpha-Aib)LDKIAQKAFVQWLIAGGPSSGAPPPS) with alpha-Aib substitutions at positions 2 and 13, as well as a 20-carbon fatty acid at Lys-20 position. Clinical trials demonstrated that tirzepatide was associated with significantly more significant weight loss than Semaglutide ^9^.

Pharmaceutical proteins, including human antithrombin III and C1-esterase inhibitor, have been successfully produced in the milk of transgenic (tg) animals ^10^. To overcome the production challenges of incretin RAs, building upon the sequences of human IgG4 Fc fragment and tirzepatide, therefore, we tested the possibility of producing Fc-fused incretin RAs in the milk of tg mice, leveraging mammary gland bioreactors for cost-effective peptide production. The construct of an optimised coding sequence driven by the beta-casein promoter was utilised to produce the tg mice, as a result, two founders efficiently secreted foreign protein in milk. Progeny exposed to tg milk displayed significantly reduced blood glucose levels and body weight, validating mammary gland bioreactors as a viable platform for producing bioactive incretin RAs.

## 2. Materials and Methods

### 2.1. Mice

Kunming mice were obtained from the laboratory animal centre of Yangzhou University and fed ad libitum standard diet containing 10% Kcal % fat. The animal study protocol was approved by the Ethics Committee of Yangzhou University (No. 202303004).

### 2.2. Vector construction

Based on the peptide sequence of tirzepatide and human IgG4 Fc fragment, the nucleotide sequence encoding GLP-1/GIP-3*(GGGGS)-Fc fragment was codon-optimised and synthesised. The expression cassette was generated after insertion into the *Xho* I site of the pBC1 milk expression vector (Invitrogen). In this construct, the coding sequence lay between exon 2 and 7 of the goat beta-casein gene, which were controlled by two tandem copies of chicken beta-globin insulator and goat beta-casein promoter (Fig 1A). The purified fragments (∼16.7 kb) digested by *Sal* I and *Not* I enzymes (New England Biolabs, Inc.) were used for pronuclear microinjection at a final concentration of 5-10 ng/μl as per the standard procedure.

**Figure 1.**
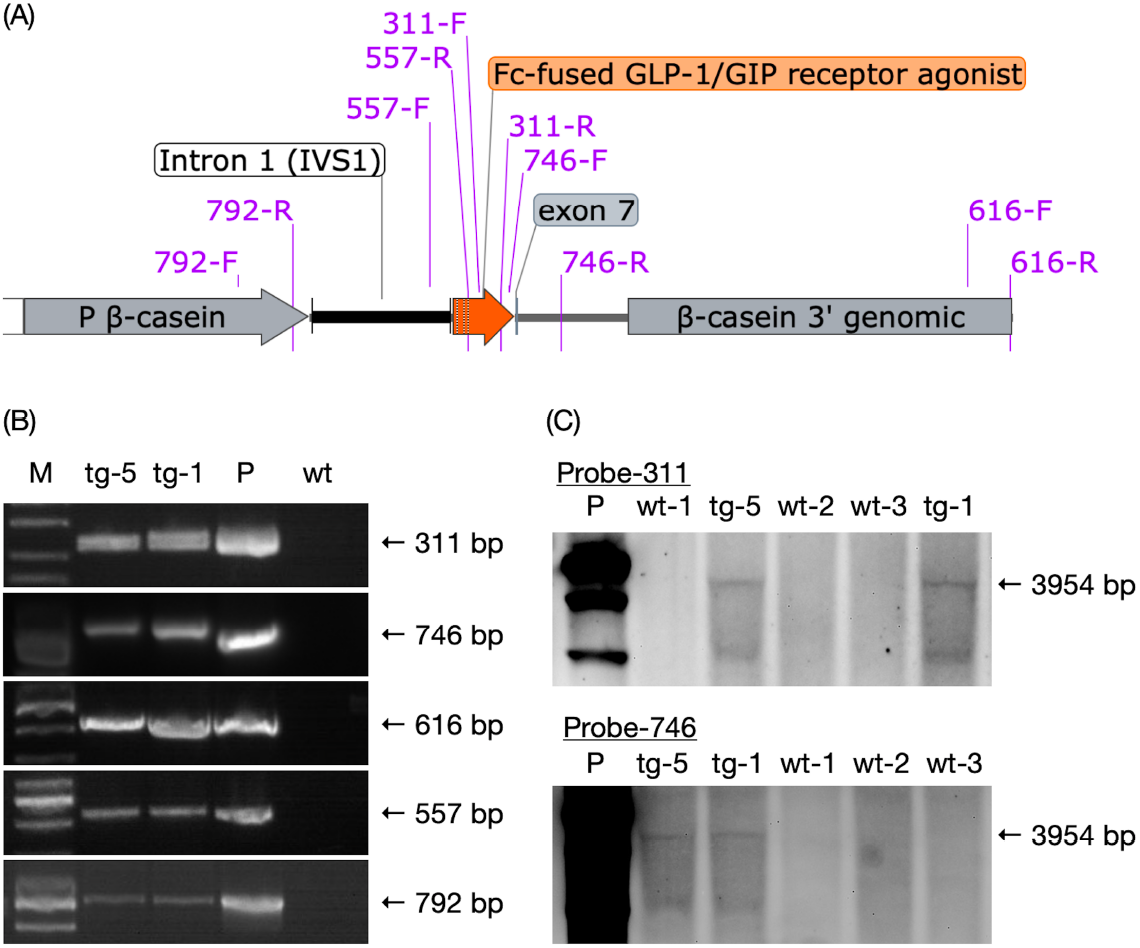
The construct illustration and molecular analysis of transgene. The optimised coding sequence of Fc-fused GLP-1/GIP receptor agonists is driven by the 2X chicken beta-globin insulator and goat beta-casein promoter, and it ended with the beta-casein 3’ genomic fragment. Positions of five primer pairs are shown in purple letters (A). PCR (B) and Southern blot (C) analysis of the genomic DNA samples from transgenic (tg), wild-type (wt) mice, and wt DNA mixed with the tg plasmid (P) indicate that tg-1 and tg-5 mice are transgenic. The sizes (bp) of aimed bands are listed on the right, and M lane represents the DNA marker.

### 2.3. Transgene analysis

Genomic DNA extracted from the tail tissues was used for nucleic acid analysis. The presence of the transgene was assayed by five primer pairs (792F: 5-AGACCCTCCTCCTGTATGGG-3, 792R: GGATGCACGGAAGTTTTGGC-3; 557F: 5-AATGGCTGGCAGTGAAAC-3, 557R: 5- CGTACTTGCTCTCGGCTGAT-3; 311F: 5-TTCAACTGGTACGTGGACGG-3, 311R: 5- GCGATGTCGCTGGGATAGAA-3; 746F: 5-AGGGGAACGTCTTCTCATGC-3, 746R: 5- AAAAGTGAGGAGGGGGCATTC-3; 616F: 5-CCCTTTCGCAGTTTGGAACC-3, 616R: 5- TGCAGGTCAACGGATCACTC-3). The PCR amplification was performed with 32 cycles at 94°C for 15 sec, 60°C for 15 sec, and 72°C for 15 sec/kb.

Southern blot was performed under standard procedure. In brief, 10 μg of genomic DNA was digested by the *EcoR* I enzyme for 12-16 h at 37°C, subjected to 1% agarose gel electrophoresis, and subsequently transferred onto the Hybond-N^+^ membrane (Amersham, UK). The membrane was hybridised with the DIG (Roche, Germany) labelled 311- or 746-bp probes according to the manufacturer’s protocol. An alkaline phosphatase-labelled anti-Dig antibody (Roche) and CDP-STAR solution were used to develop the photos.

### 2.4. Milk protein analysis

Milk was harvested on Day 5, 8, 10, 14, or 21 from oxytocin-treated tg or wild-type (wt) mothers (2-3 i.u. per mouse) postpartum (Day 0) according to the standard procedure for immediate analysis or stored at -80°C for later analysis.

Western blot was performed according to the standard procedures. In brief, 5 µl of 1:20 diluted whole milk on Day 8 or Day 10, or 20 µg protein extracts of each tissue from a tg and a wt offspring of tg-1 at 10 days of age, were separated by 10% SDS-PAGE gel and transferred to the PVDF membranes (Millipore ISEQ00010, Merck). The blocked membrane was hybridised with the anti-tirzepatide polyclonal antibody (Biolab Reagents, Wuhan, China; PHK13902) or beta-Actin (Proteintech Group, Inc., China; 20536-1-AP) followed by HPR-labelled goat anti-rabbit IgG (Santa Cruz Biotechnology; sc-2004); or the membrane was directly hybridised with HPR-labelled goat anti-human IgG Fc antibody (ThermoFisher Scientific Inc, Shanghai, China; A18817). The chemiluminescent signal was developed using the SuperSignal™ West Femto substrate (ThermoFisher). The prestained protein marker was obtained from Vazyme Ltd. (Nanjing, China; MP102).

ELISA was performed to detect the targeted protein concentration according to the standard procedures. Briefly, 100 µl of 1:20 diluted whole milk was quantified using the HRP-labelled goat anti-human IgG Fc secondary antibody (ThermoFisher, A18817). The human IgG (Beyotime Biotechnol., Shanghai, China; A7001) was the standard sample.

### 2.5. Measurements of blood glucose level and body weight of F1 offspring

Within the group of tg-1, the female founder and one of its non-tg littermates got mated with the same wt-A male on the same day, and each of them separately delivered and fostered babies by itself for the remaining period.

Within the group of tg-5, the female founder and one of its non-tg littermates got mated with the same wt-B male on the same day, and two mothers resumed fostering their babies together within the cage. On Day 8, the two mothers and 22 pups were randomly and equally divided into individual cages to continue breastfeeding. Fresh blood from the tail tips of non-fasted F1 pups was used to measure circulating glucose levels using a handheld glucose monitor (Accu-Chek^®^ Performa Blood Glucose Meter, Roche); meanwhile, the body weight of each pup was measured.

### 2.6. Statistical analysis

Data are presented as mean ± standard deviation (SD). A comparison between wt and tg samples was performed using an unpaired two-tailed t-test with Welch’s correction where appropriate. Analysis was conducted using GraphPad Prism 8.0 (GraphPad Software Inc.; La Jolla, CA, USA), and p < 0.05 was considered statistically significant.

## 3. Results

### 3.1. Expression of Fc-fused GLP-1/GIP receptor agonists in the mammary gland

Two tg female founders, namely tg-1 and tg-5 from the cohort of 95 mice derived from injected embryos, were confirmed by analyses of PCR (Fig 1B), DNA sequencing, and Southern blot (Fig. 1C). Milk samples from lactating founders were analysed via Western blot (anti-tirzepatide and anti-Fc antibodies). Both tg-1 and tg-5 expressed Fc-fused GLP-1/GIP RAs (∼37 kDa) exclusively in milk, with no detection in somatic tissues such as heart, liver, spleen, lung, kidney, and brown adipose tissues (Fig. 2). ELISA quantified that the concentrations of Fc-fused protein were 0.8 and 1.33 g/l in the tg-1 milk on Day 5 and Day 10; as well as 1.11, 1.15, and 1.42 g/l in the tg-5 milk on Day 8, Day 10, and Day 14 respectively. The results suggest that the Fc-fused protein has been efficiently produced in the milk of tg animals.

**Figure 2.**
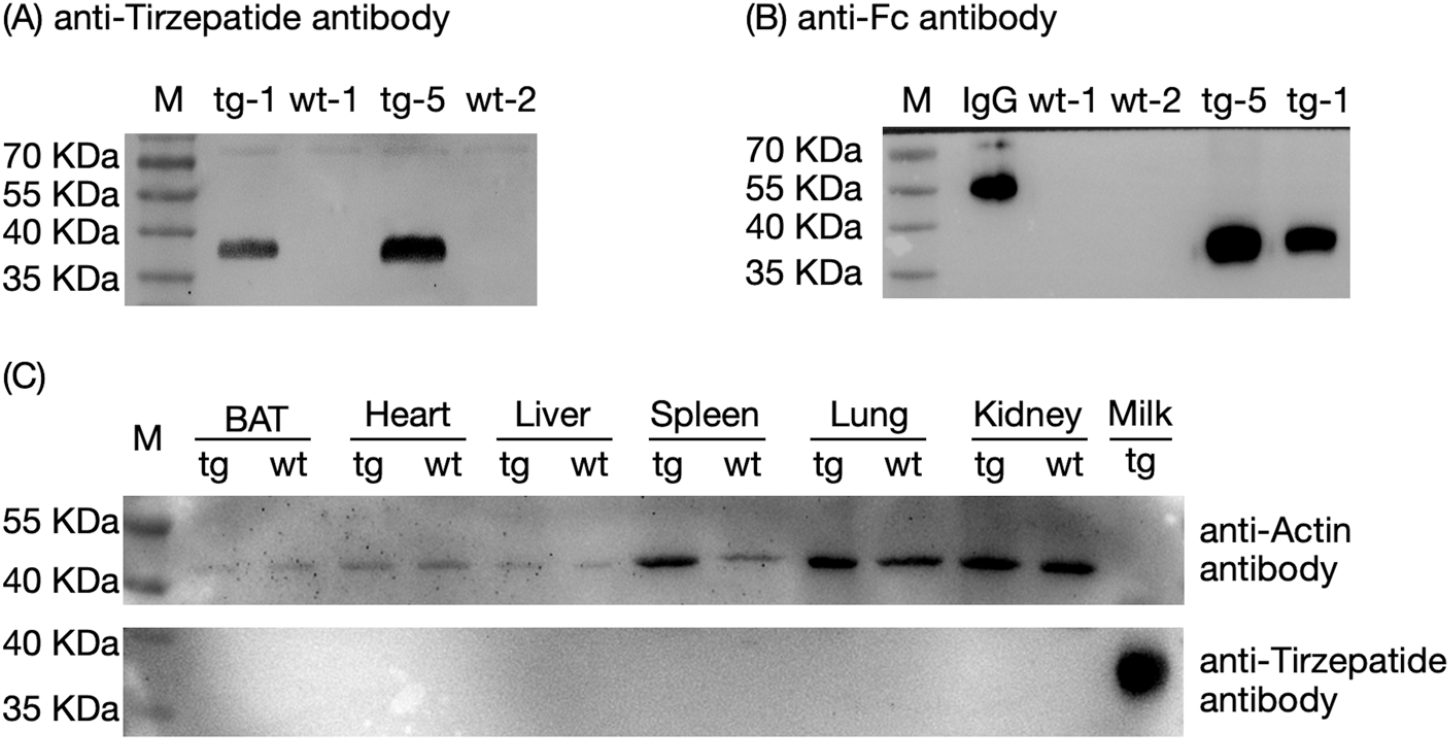
Western blot analysis of Fc-fused GLP-1/GIP receptor agonists in the milk of transgenic (tg) mice. Western blot analyses revealed that Fc-fused GLP-1/GIP receptor agonists (∼37 kDa) exist only in tg-1 and tg-5 milk using anti-tirzepatide (A) or anti-Fc antibodies (B), but not other tg tissues including brown adipose tissue (BAT), heart, liver, spleen, lung and kidney, as well as wild-type (wt) tissues (C). Milk samples were collected on Day 8 (A, B) or Day 10 (C), and tissues came from the Day-10 F1 offspring of the tg-1 founder (C). Human IgG (heavy chain ∼55 kDa) is the positive sample (B), and M represents the protein marker.

### 3.2. Dual glucose-lowering and weight-reducing effects of the tg milk on fostered pups

Both tg founders exhibited normal reproductive capacity and delivered their babies successfully. Blood glucose levels and body weight of their F1 pups were further assessed to evaluate the biological activity of the Fc-fused GLP-1/GIP RAs in the milk of tg mice. Tail-tip blood glucose levels in offspring at 8, 10, 14, and 21 days of age revealed that pups from the tg-1 founder exhibited consistently lower glucose levels compared to age-matched wt controls, with reductions of 10.37% (p < 0.05), 20.92% (p < 0.01), 10.88%, and 16.97% (p < 0.001), respectively (Fig. 3A).

**Figure 3.**
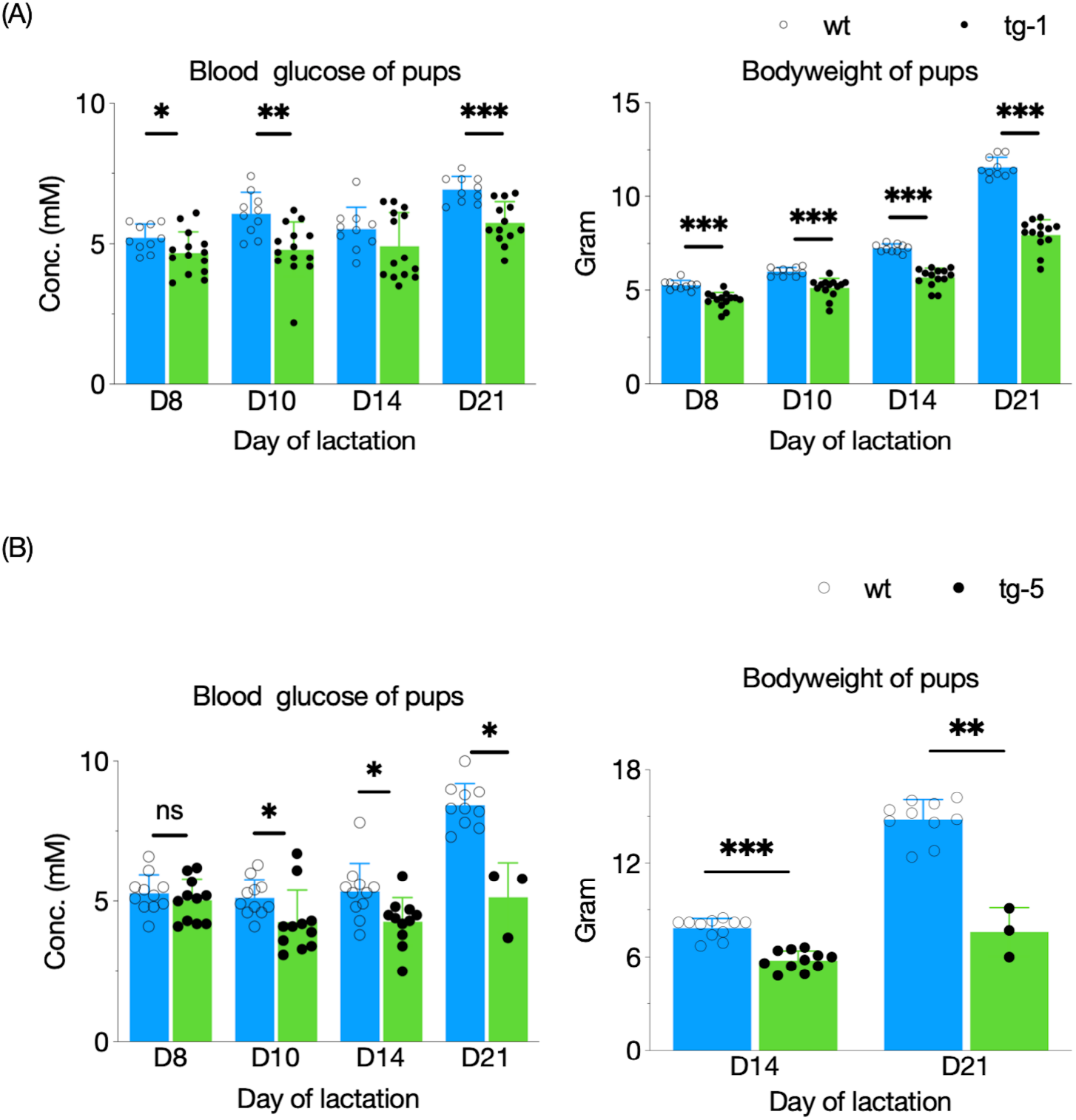
Decrease of blood glucose levels and body weights of offspring fostered by the tg-1 (A) or tg-5 (B) mice. Bars represent the mean ± SD. * indicates p < 0.05; ** indicates p < 0.01; and *** indicates p < 0.001.

Concurrently, the body weights of tg-1 offsprings at these ages were markedly lower (p < 0.001) than wt counterparts, showing reductions of 14.86%, 14.36%, 21.88%, and 31.25%, respectively; however, all F1 pups of the tg-1 founder exhibited growth until Day 21.

In the tg-5 group, despite being co-nursed by tg-5 and wt mothers until Day 8, offspring continuously nursed by the tg-5 founder showed 16.87%, 19.93%, and 39.39% blood glucose reduction on Day 10, 14, and 21 compared to wt controls (p < 0.05; Fig. 3B). Concurrently, pups fostered by the tg-5 founder exhibited a 26.42% reduction (p < 0.001) in body weight by Day 14. Notably, severe mortality occurred in the tg-5 group on Days 16–18, with 8 out of 11 pups dying; only three survived to Day 21, displaying a dramatic 48.61% weight reduction (p < 0.01) compared to wt-nursed offspring that all recovered glucose levels and grown normally (Fig. 3B). These findings indicate that the Fc-fused GLP-1/GIP RAs in the tg milk possesses both hypoglycaemic and weight-suppressing effects.

## 4. Discussion

Previous studies indicated that beta-casein promoter could efficiently drive the foreign gene to express recombinant proteins such as human alpha-1-antitrypsin as high as 35 g/L in the milk of tg animals ^11^ and has been successfully utilised to produce mammary bioreactor models until now ^12^. With the success of early tg therapeutic proteins, including human antithrombin III, C1esterase inhibitor, and alpha-1 antitrypsin, it is clear that the pharmaceutical and biotechnology industries have accepted the production of valuable therapeutic proteins in the milk of tg animals.

The neonatal Fc receptor can mediate the intestinal absorption of IgG (Paveglio et al., 2012), prompting us to directly evaluate the biological activities of recombinant milk protein by measuring the blood glucose levels and body weights of pups nursed by tg founders. In addition to the fact that tg-1 and tg-5 founders specifically expressed GLP-1/GIP RAs in the mammary gland rather than in other tested tissues, there was no significant difference in blood glucose levels or body weights between eight tg and six non-tg littermates produced by tg-1 mice from Day 10 to Day 21 (data not shown). Similarly, there was no difference in blood glucose levels between four tg and seven non-tg pups nursed by the wild-type mother in the tg-5 group. These results suggest that foreign incretin RAs are exclusively expressed in the mammary gland and are safe for the host animals.

The receptor-binding sequence HAEGTFTSDVSSYLEGQAAKEFIAWLVKGR (residues 7-37) of the original GLP-1 protein can be rapidly degraded primarily by the dipeptidyl peptidase-4, resulting in a half-life of approximately 2 minutes. This proteolytic enzyme cleaves the peptide bond between Ala-8 and Glu-9. To overcome this, the replacement of Ala-8 by Gly-8 or α-aminoisobutyric acid is usually performed in therapeutic proteins, including Semaglutide and Tirzepatide. To avoid serious harm to pups, the residues Ala-8 and Glu-9 in the original position were preserved intact in this study; even so, the progeny exposed to tg milk exhibited drastic metabolic effects, especially tg-5-nursed pups experienced high mortality post-Day 16, likely due to overdosing. It indicates that the Fc fragment does extend the half-life of RAs.

This study demonstrates the feasibility and efficiency of producing bioactive Fc-fused GLP-1/GIP RAs in the milk of tg animals, aligning with prior bioreactor models for pharmaceuticals ^10^, and the Fc-fusion strategy mirrors clinical successes like dulaglutide ^5^. Although the ability of tg production to compete with chemical synthesis or conventional cell culture production will remain a challenge, this platform could revolutionise the cost-effective and supplemental production of incretin therapies. Thus, we anticipate the eventual production of large-scale, high-level incretin RAs using larger animals.

## Abbreviations

BAT: brown adipose tissue
GIP: glucose-dependent insulinotropic polypeptide
GLP-1: glucagon-like peptide-1
RAs: receptor agonists
tg: transgenic
wt: wild-type

## Ethics approval and consent to participate

All animal experiments were performed following the Committee for Experimental Animals of Yangzhou University.

## Institutional Review Board Statement

The animal study protocol was approved by the Ethics Committee of Yangzhou University (protocol code No. 202303004 and dated 10 March 2023).

## Consent for publication

Not applicable

## Availability of data and materials

The datasets used and analysed during the current study are available from the corresponding author upon reasonable request.

## Authors’ contributions

Conceptualization, K Gou; Data curation and Investigation, Y Rao, B Wang, S Yu; Formal analysis, Y Rao and K Gou; Methodology, Validation and Visualization, Y Rao, S Yu; Project administration, Supervision and Writing – review & editing, S Cui and K Gou; Writing – original draft, Yu Rao; All authors proofread, made comments, and approved the report.

## Acknowledgements

We thank Ms Wen-Jing Gou’s language help in preparing this manuscript.

